# Lineage segregation in human pre-implantation embryos is specified by YAP1 and TEAD1

**DOI:** 10.1101/2022.09.29.509946

**Authors:** Marius Regin, Wafaa Essahib, Andrej Demtschenko, Delphine Dewandre, Laurent David, Claudia Gerri, Kathy Niakan, Greta Verheyen, Herman Tournaye, Johan Sterckx, Karen Sermon, Hilde Van de Velde

## Abstract

We know that polarity and YAP1 play a key role in trophectoderm initiation in compacted human embryos, however we know little about the TEAD family of transcription factors that become activated by YAP1 and especially if they play a role during epiblast and primitive endoderm formation. Here we show that compaction occurs heterogeneously between the 8- and 16-cell stages. While 8-cell stage blastomeres are not yet polarized, polarized outer cells and non-polarized inner cells arise in compacted 16-cell stage embryos. While trophectoderm specifiers TEAD1, YAP1 and GATA3 mostly co-localise in the nuclei of polarized outer/trophectoderm cells, they are also found in some cells of compacting embryos before polarity is established indicating that differentiation into trophectoderm cells can be initiated independently of polarity. In the inner cell mass, TEAD1 and YAP1 also distinguish GATA4 positive cells in a salt-and-pepper distribution and in the sorted primitive endoderm cells. Our detailed roadmap on polarization, compaction, position and lineage segregation events during human preimplantation development paves the road for further functional studies. Fundamental knowledge of lineage segregation events will eventually explain how and why embryos fail to develop further before or during implantation.

## Introduction

During human pre-implantation development a totipotent zygote develops into a competent blastocyst ready for implantation 7 days post fertilisation (dpf). The segregation into trophectoderm (TE) and the pluripotent inner cell mass (ICM) cells represents the first lineage differentiation event. The ICM cells further segregate into epiblast (EPI) and primitive endoderm (PrE) cells, which represents the second lineage differentiation event. TE cells are responsible for implantation while PrE cells develop into the yolk sac and both are precursors of embryonic parts of the placenta (chorion); the EPI cells give rise to the embryo proper.

Although extensively studied in the mouse model, the molecular events driving lineage segregation to either TE, EPI or PrE cells in human embryos remain largely unknown. In the mouse, the Hippo signalling pathway plays a crucial role in both TE and EPI differentiation, with the TEA-domain transcription factor family (TEAD1-4) and co-activator Yes-associated-protein 1 (YAP1) as key components (Nishioka *et al*., 2008, 2009; Ralston *et al*., 2010; Hirate *et al*., 2013; Hashimoto and Sasaki, 2019). Differences in cell polarity model are thought to drive the first lineage differentiation event in the mouse. Mouse embryos compact at the 8-cell stage with all blastomeres being polarized (Levy *et al*., 1986). Polarity is marked by the presence of an apical cap composed of the PAR complex proteins (PAR3, PAR6 and atypical Protein Kinase C (aPKC)) and p-ERM, which is a component of the microvilli linking F-actin to the apical membrane (Plusa *et al*., 2005; Alarcon, 2010; Zhu *et al*., 2017). During the 8-to 16-cell stage divisions, a symmetric division through the meridian yields two daughter cells with a polar cap that are located in an outer position. After an asymmetric division through the equator, cells that inherit the polar cap will remain in an outer position whereas non-polarized cells will be relocated towards an inner position (Johnson and Ziomek, 1981; Anani *et al*., 2014; Lorthongpanich and Issaragrisil, 2015). This is crucial for establishing an inner/non-polarized and outer/polarized cellular axis in the mouse embryo. Non-polarized inner cells exclude YAP1 from the nuclei through phosphorylation of serine127 by the large tumour suppressor protein kinase (LATS1/2). Consequently, *Cdx2* expression is downregulated, whereas SOX2 protein levels increase maintaining pluripotent inner cells and rendering them precursors of the ICM (Wicklow et al., 2014). In contrast, in polarized outer cells LATS1/2 activity is restricted by the presence of the polar region and TEAD4/YAP1 nuclear activity leads to the upregulation of *Cdx2* and *Gata3* expression (Nishioka *et al*., 2008, 2009; Ralston *et al*., 2010; Hirate *et al*., 2013). The first lineage segregation event is also regulated by other mechanisms such as cortical tension (Maître *et al*., 2015; Maître, 2017; Zhu *et al*., 2017).

In the mouse, the second lineage segregation event is regulated by the FGF4/FGFR2 signalling pathway (Chazaud *et al*., 2006; Yamanaka *et al*., 2010). In the ICM cells, the NANOG positive precursor EPI cells secrete FGF4 for which the GATA6 positive precursor PrE cells have FGFR2 receptors. The EPI and PrE cells first display a salt-and-pepper pattern whereafter the cells are sorted and an inner population of EPI cells are covered by a monolayer of PrE cells facing the cavity just before implantation. Recently TEAD1/YAP1 signalling has also been found to play a role in the establishment of the EPI cells (Hashimoto and Sasaki, 2019), but it is not known how the two signalling pathways act together.

Recent single cell transcriptomic analysis of human embryos gave also insight into transcription factors that are associated with each lineage differentiation event during preimplantation development (Yan *et al*., 2013; Blakeley *et al*., 2015; Petropoulos, 2016; Stirparo *et al*., 2018; Meistermann *et al*., 2021; Radley *et al*., 2022). We and others demonstrated by careful embryo staging using morphokinetic analysis that human embryos initiate compaction at variable times between the 8- and the 16-cell stage (Gerri *et al*., 2020). The first lineage differentiation in compacted human embryos is orchestrated by polarity and involves YAP1 and GATA3 signalling (Gerri *et al*., 2020) and phospholipase C signalling (Zhu *et al*., 2021). Few data are available on the second lineage differentiation in human blastocysts, but it appears independent of FGF signalling (Kuijk *et al*., 2012; Roode *et al*., 2012). Based on single-cell transcriptomics data, competing hypotheses on the second lineage differentiation in human blastocysts have been proposed. Petropoulos et al. suggested that EPI and PrE lineages arise simultaneously with TE cells on 5 dpf (one step model) (Petropoulos, 2016) whereas Meistermann et al. proposed that the first and the second lineage differentiation events appear consecutively (two step model) (Meistermann *et al*., 2021) as in mice (Chazaud and Yamanaka, 2016). It is also unclear whether ICM cells segregate into EPI and PrE cells (Radley *et al*., 2022) as in mice (Chazaud and Yamanaka, 2016) or whether PrE cells differentiate from the EPI cells on 6 dpf (Meistermann *et al*., 2021). This lack of clarity may have been caused by the heterogeneous development of outbred human embryos *in vitro*, different laboratory culture conditions and the lack of systematic coupling of morphology with the underlying analysis. Moreover, limited data are available on the transcription factors and signalling pathways at the protein level and very few functional studies have been performed.

Our current aim was to provide a comprehensive spatial and temporal map of the processes of compaction, polarisation, position and blastulation, and the first and second lineage differentiation events in human pre-implantation embryos. Through the investigation of the main proteins in well-defined human pre-implantation stages we demonstrate a human-specific chain of events during compaction and polarisation in outer and inner cells. We also reveal that in (precursor) TE and PrE cells of human pre-implantation embryos the transcription factor TEAD1 and co-activator YAP1 co-localize.

## Material and Methods

### Ethical statement

This project was approved by the Local Ethical Committee of the UZ Brussel (BUN 143201526417) and the Belgian Federal Committee for research on human embryos (AdV057). We used cryopreserved human embryos donated to research from IVF-ICSI patients after informed consent and the expiration of the 5-year cryopreservation duration. Mature oocytes were obtained from egg bank donors who voluntary donated gametes for research.

### Embryo warming and culture

Vitrified high quality 8-cell stage human embryos (3 dpf) were warmed (Vit Kit -Thaw, Irvine Scientific, USA) and cultured until 6 dpf in individual microdroplets of 25µl medium (Quinn’ s Advantage™ Blastocyst media, Origio, USA) under a layer of paraffin oil (OVOIL™ - culture oil, Vitrolife, USA) to prevent evaporation of medium. Embryos were incubated at 37°C, 5% O2 and 6% CO2 tension. Warmed embryos were scored for viability per blastomere after warming, overall survival of the embryos was at 99%. Embryos with signs of degeneration in more than one cell were excluded as well as embryos that showed more than 10% fragmentation, with irregular size of blastomeres, increased amounts of vacuoles or granulation and arrested embryos.

### Developmental timeline of the human embryo

The embryos were scored according to their developmental stage displaying morphological key differences rather than defining them according to their chronological age (days post fertilisation or dpf) (Supplementary Table 1). The compaction process was divided into three subgroups. At the beginning of compaction (C0) the outlines of the individual blastomeres within the embryo were clearly visible. However, the cell shape appeared slightly elongated. Shortly after, the cellular borders began to fade as a result of increased intercellular contacts due to adherence junctions (C1). At the end of compaction (C2) the blastomeres had maximized their contact sites and the embryo appeared as a sphere of contour-less tightly packed cells. Inner cells were identified as cells that are entirely separated from the perivitelline space and were therefore enclosed by cellular contacts on all sides. The blastulation process was divided into four subgroups. Transition from the fully compacted embryo to the early blastocyst stage (B0) was marked by a slight decompaction as cellular outlines were once again visible, small cavities were formed and sickle-cell shaped outer cells became apparent. Subsequently one cavity was formed (B1). As soon as the ICM was visible, the outer cells were referred to as TE in a full blastocyst (B2). As fluid accumulated further the embryo started to expand as an increase in size could be observed due to TE cell proliferation and ZP thinning (B3). Subsequently the blastocyst expanded (B4) and started to hatch out of the ZP (B5) until it was fully hatched (B6).

### Immunofluorescence

Cultured embryos were fixed in 4% PFA/PBS for 10 minutes at room temperature (RT) to crosslink proteins and preserve cellular integrity. After fixation embryos were washed three times for 10 minutes in 2% BSA/PBS before permeabilization of membranes in 0,1% TritonX/PBS for 20 minutes at RT to increase access for antibody binding. Subsequently embryos were placed in 10% FBS/PBS blocking solution for 1h at RT to decrease unspecific antibody binding before incubation with primary antibody or control-IgG antibody (Supplementary Table 2) in 2% FBS/BSA overnight at 4°C. Embryos were washed in 2% BSA/PBS before incubation with secondary antibody (Supplementary Table 2) at 10 µg/ml for 2h at RT in the dark, after 1h of secondary antibody incubation fluorochrome conjugated phalloidin (Alexa Fluor™ 546 Phalloidin, Invitrogen™) was added to the secondary antibody solution for F-actin membrane staining for another hour. Embryos were washed again in 2% BSA/PBS before incubation with Hoechst 33342 (Molecular probes®, Life Technologies) for nuclear staining. Eventually embryos were mounted in 1xPBS on poly -L-lysine coated glass slides (Poly-Prep Slides, Sigma-Aldrich) to prevent collapse and movement during visualization. Round coverslips were sandwiched between the glass slide and the coverslip (Corning® Cover Glasses, #1.5, Sigma-Aldrich) and glued together with nail polish to prevent squeezing of the embryo.

### Confocal microscopy and visualisation

Fluorescence of immunostained embryos was detected with a confocal microscope (LSM800 equipped with Airyscan, ZEISS). Whole embryo z-stack scans were carried out to reconstruct the 3D anatomy of the embryo. To omit spectral overlap between fluorophores, emission filters for each fluorophore were employed. Z-stacks were processed with the Zen-Blue software (ZEISS) and image sequences were exported and analysed for cell counts with ImageJ and Arivis Vision 4D (3.4.0). The number of nuclei (positive for DNA) was assessed using the Blob finder function of Arivis. All other counts were assessed manually. We considered a nucleus positive for a protein when the signal visibly exceeded the background and/or when the signal was clearly delineated from the cytoplasm. We confirmed our manual approach by measuring fluorescence intensities of YAP1 and GATA3 at the earlier stages when signals started to emerge, thus when weak signals are more prone to operator error (Fig. 4B) using the 3D Fiji/Image J Suite as previously described (Gerri *et al*., 2020, 2022).

### Statistics

We used a two tailed Mann Whitney U test for the data regarding the fluorescence intensities (Fig. 4B). All graphs and statistics were generated using GraphPad Prism (9.0.0.).

## Results

### Inner and outer cells are established in the compacted human embryo

In order to establish the number of cells at each embryonic developmental stage as described in the clinical grading system, we first analysed the appearance and the number of inner and outer cells in relation to the total number of cells by confocal microscopy (Fig. 1). Cell counts were determined by analysing confocal Z-stack scans for nuclear Hoechst and membrane F-actin staining (Fig. 1A).

**Figure 1:**
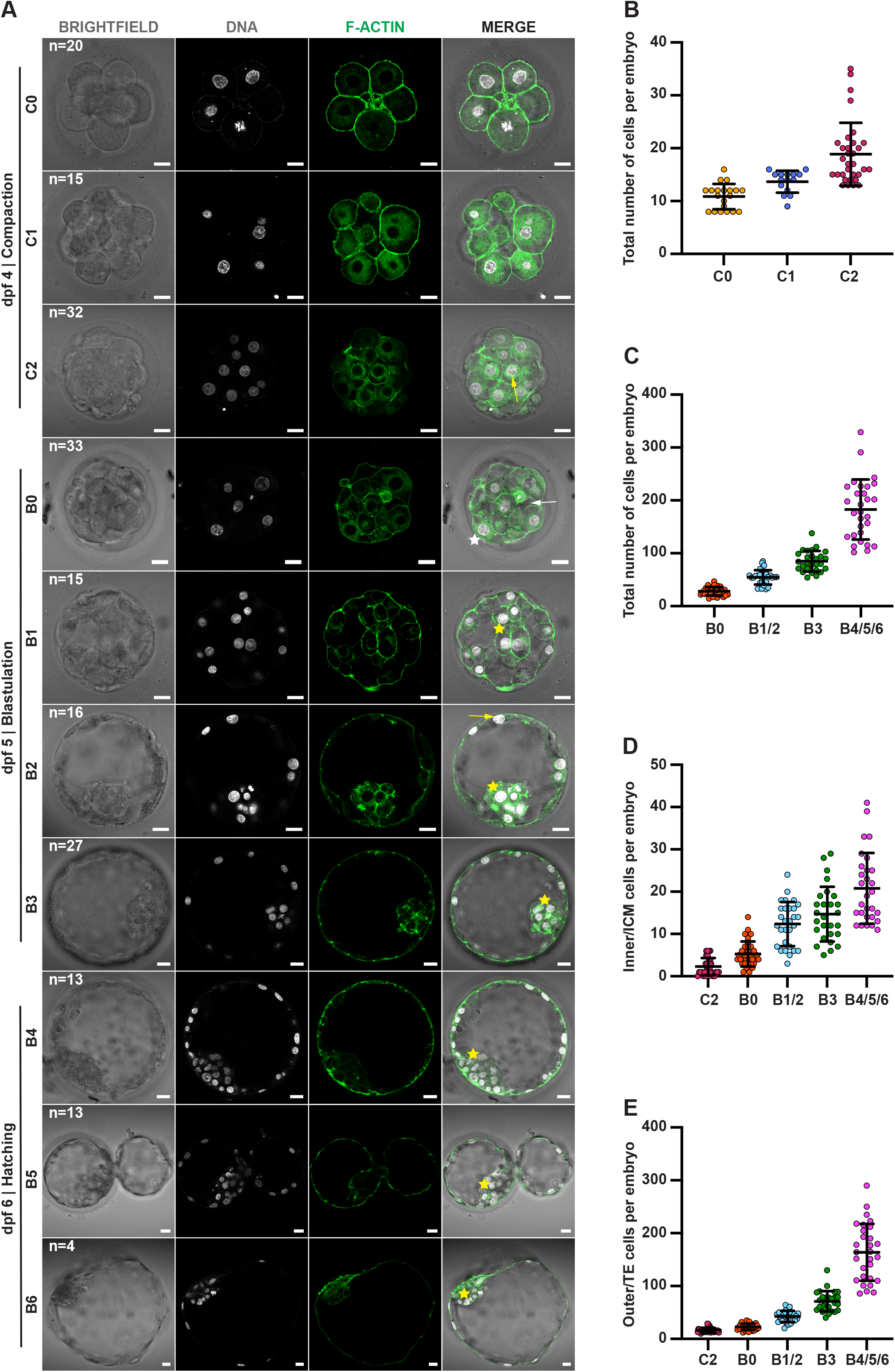
Inner and outer cells are established in the compacted human embryo. (A) Developmental timeline of the human embryo based on morphology and distribution of F-actin (green) and DNA (white) at 4 days post fertilisation (dpf) (C0-C2), 5 dpf (B0-B3) and 6 dpf (B4-B6). At the beginning of compaction (C0) the contact between de blastomeres increases and the cell shape appears slightly elongated, but the outlines are still clearly visible. Shortly after, the cellular borders begin to fade as a result of increased intercellular contacts due to adherence junctions and the embryo is compacting (C1). When compaction is completed (C2) the blastomeres have maximized their contact sites and the embryo appears as a sphere of contour-less tightly packed cells. During this stage the first inner cells (yellow arrow) appear. The initiation of blastulation (B0) is marked by a slight decompaction (white arrow) as cellular outlines are visible and the outer cells become sickle-cell shaped (white star) with a cavity (B1). As soon as the ICM (B2-B6, yellow stars) is visible, the outer cells become trophectoderm (TE) cells (yellow arrow). During the B3 stage, the blastocyst expands with coinciding zona-pellucida (ZP) thinning. At 6 dpf an increase in blastocyst size can be observed due to TE-cell proliferation (B4). Subsequently the blastocyst starts to hatch (B5) until fully hatched out of the ZP (B6). (B) Total number of cells per embryo during compaction (C0-C2) and (C) blastulation (B0-B6) (D) Inner/ICM cells per embryo from C2-B6. (E) Outer/TE cells per embryo from C2-B6. Brightfield images were taken after fixation. Scale bars, 20µm. All error bars are mean ± SD.

We observed no inner cells, defined as cells completely separated from the perivitelline space, during the compaction process (Fig. 1D, Fig. 2A). At the onset of compaction (C0), the embryos contained 10.9±2.5 cells and compacting C1 embryos contained 13.6±2.1 cells (Fig. 1B). The first inner cells were observed in the fully compacted C2 embryos, which on average contained a total number of 18.9±6.1 cells (Fig. 1B), and showed a range of inner cells between 0 (5/30 embryos) and 6 (Fig. 1D), with a mean of 2.3±0.7, while 16.8±4.7 cells were outer cells (Fig. 1E).

**Figure 2:**
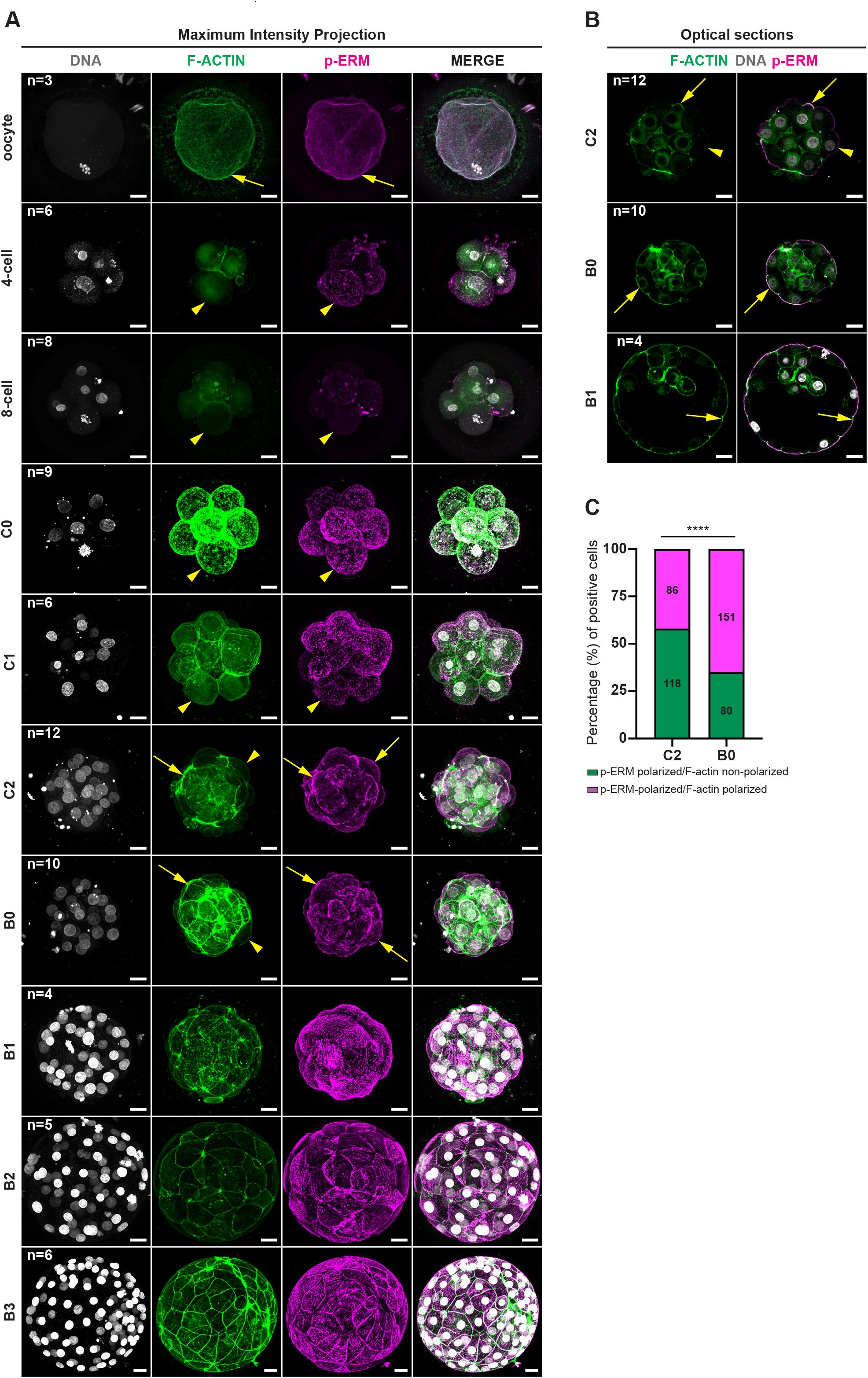
P-ERM apical polarity proceeds F-actin apical polarity in the compacted human embryo. (A) Maximum intensity projections of different developmental stages analysed for DNA (white), F-actin (green) and p-ERM (magenta). Presence of F-actin and p-ERM apical polarity (yellow arrows); absence of apical polarity (yellow arrow heads). (B) Optical section of embryos (C2, B0, B1) analysed for F-actin (green), DNA (white) and p-ERM (magenta). F-actin apical polarity indicated by yellow arrow, absence indicated by yellow arrowheads. (C) Percentage of positive cells at the C2 and B0 stage that show either p-ERM and F-actin polarisation or p-ERM polarisation without F-actin (****p<0.00001, Fisher-exact-test). Brightfield images were taken before fixation. Scale bars, 20µm.

When small cavities appeared, the outer cells changed morphologically into sickle-shaped cells (Fig. 1A, B0, white star). The total number of cells increased from an average of 28.1±7.8 cells at the onset of blastulation (B0) to 54.8±13.7 in full blastocysts (B1/2) and 81.4±14.8 (B3) upon expansion at 5 dpf and to 155.4±39.5 in fully expanded blastocysts (B4/5/6) at 6 dpf (Fig. 1C). By comparing the cell counts for inner/ICM and outer/TE cells throughout development, we observed that while the number of inner/ICM cells followed a linear increase, the number of outer/TE cells increased exponentially (Fig. 1D-E). We conclude that compaction occurs heterogeneously, i.e. without apparent established course of events, between the 8- and 16-cell stages and inner cells first appear in compacted human embryos.

### Polarity marker p-ERM is apically localised in outer cells of compacted human embryos and precedes F-actin

We aimed to identify differences in apical polarity between inner and outer cells throughout human pre-implantation development by co-staining for F-actin and p-ERM. F-actin has already been used as an apical marker in human embryos by Zhu et al. (Zhu *et al*., 2021); p-ERM has not yet been investigated in human embryos but is well-described in the mouse model (Anani *et al*., 2014; Zhu *et al*., 2020).

First, we determined the spatial and temporal localisation of F-actin during human pre-implantation development (Fig. 1A, Fig. 2A-B). While in oocytes and cleavage-stage embryos F-actin was mainly diffuse in the cytoplasm, it became concentrated at the cortex of all the cells at the start of compaction (C0) to remain there during the compaction process (C1). F-actin started to appear concentrated in the apical region of outer cells when compaction was completed (C2) and it gradually increased at the start of cavitation in B0 blastocysts; in C2 and B0 stages F-actin also appeared as a ring around the nucleus. In B1 blastocysts F-actin was apically polarized in all the outer cells. From the B1 blastocyst stage onwards it became gradually concentrated in the intercellular junctions between the sickle-shaped outer cells. Finally, F-actin was highly concentrated in the junctions of the TE cells (B2-B6) but remained perinuclear and at the cortex in the ICM cells in full B2 and expanding B3 blastocysts (Figure 2A) and in inner cells of expanded B4 and hatching/hatched B5/B6 blastocysts (Figure 1A). We conclude that F-actin is localised apically in all the outer/TE cells from the early B1 blastocyst stage onwards.

Next, we analysed the pattern of p-ERM during pre-implantation development (Fig 2A-B). Diffuse presence of p-ERM without polarity was observed in oocytes and blastomeres of cleavage-stage embryos. P-ERM gradually became localised at the cell-contact-free surface during compaction in C0 and C1 embryos without reaching apical polarity. Full apical polarity of p-ERM was observed in all outer cells of all compacted C2 embryos, while it was absent at sites of cellular contact. P-ERM remained apical in early blastocysts (B0 and B1), then shifted from the apical cap to intercellular regions during blastocyst expansion (B2-B3). We conclude that p-ERM is localized in the apical cap of all outer cells in compacted C2 embryos and blastocysts.

Since the immunofluorescence analysis suggested that p-ERM polarity precedes F-actin polarity, we analysed the co-localisation of p-ERM and F-actin to define the precise order (Fig. 2A and Fig2C). At the compacted C2 stage all outer cells had an apical p-ERM cap, but only 42.2% of these cells had polarized F-actin, while this number rose to 65.4% of outer cells in the early B0 blastocysts that showed apical co-localisation of p-ERM and F-actin, suggesting that p-ERM polarizes before F-actin (p<0.00001, Fisher-exact-test). From the B1 blastocyst stage onwards, p-ERM and F-actin co-localized in the apical cap of all outer cells. We observed a shift of p-ERM distribution from the apical surface of the blastomeres to sites of intercellular contact coinciding with F-actin re-allocation during blastocyst expansion (B2/B3) (Fig 2A-B). We conclude that p-ERM polarisation is completed by the compacted stage in human embryos and precedes F-actin apical polarity.

### Nuclear YAP1 precedes polarity and mainly specifies polarized outer/TE cells

Because of their important role in cellular polarity and lineage segregation in the mouse, we co-stained for p-ERM and YAP1 (Fig. 3A). While few cells (19.5%) were positive for nuclear YAP1 at the start of compaction (C0), their number markedly increased (56.1%) at the compacting embryo stage (C1) in the presence of non-polarized p-ERM (Fig. 3B). At the compacted embryo stage (C2), all outer cells were polarized and most of these cells (84.6%) displayed high levels of nuclear YAP1; all inner cells were non-polarized and in 25% of these cells nuclear localization of YAP1 was found (Fig. 3C, C2 panel). In general, throughout the blastocyst stages B0-B3 on 5 dpf, all embryos showed polarized outer/TE cells positive for YAP1 and non-polarized inner/ICM cells negative for YAP1 (Fig. 3D). The outer cells of B0 embryos contained the largest number of cells that were mostly YAP1 positive and polarized (99%) with the exception of one non-polarized cell. The majority of inner cells lacked YAP1 and polarity (84%) but in rare cases they were YAP1 positive and non-polarized (4%) or polarized (12%) (Fig. 3C, Fig. S1A, first row). At the B1-B3 stage, we found that 97.1% of the outer/TE cells were polarized and YAP1 positive in B1, 94.8% in B2 and 95% in B3. Most non-polarized inner/ICM cells lacked YAP1: 100% at B1, 97.6% at B2 (with the exception of one cell that was polarized and YAP1 positive) and 100% at B3 (Fig. 3D). Throughout the blastocyst stages, we also found a small population of polarized TE cells negative for YAP1 (3.4%) (Fig. S1A, second row). We conclude that nuclear YAP1 may precede the establishment of the apical polar cap and becomes mainly restricted to polarized outer/TE cells from the compacted embryo stage onwards.

**Figure 3:**
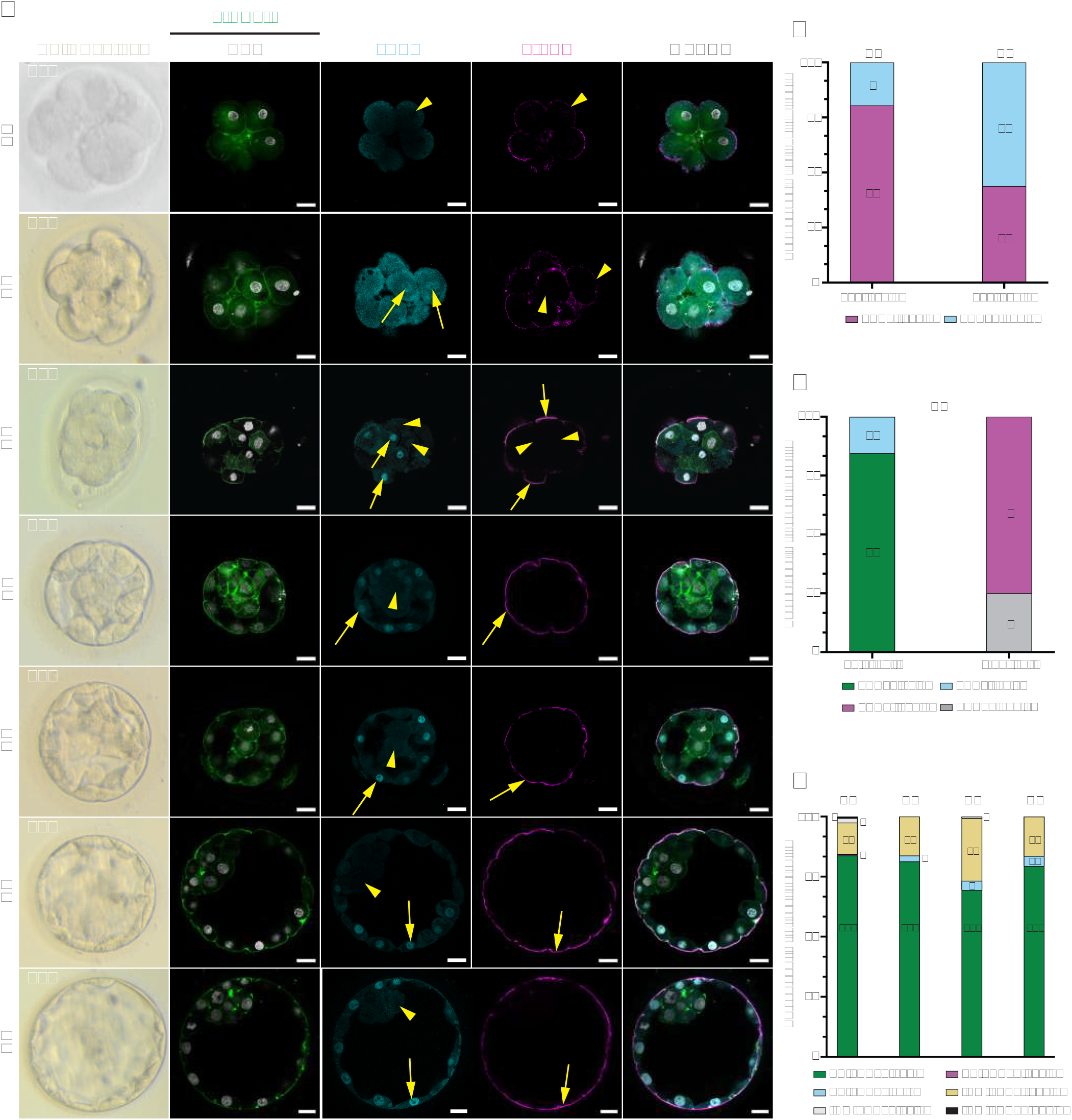
YAP1 expression becomes mainly restricted to polar outer cells from the compacted human embryo stage onwards. (A) Optical sections of embryos (C0-B3) analysed for DNA (white), F-actin (green), YAP1 (turquoise) and p-ERM (magenta), presence indicated by yellow arrow, absence indicated by yellow arrowheads. (B) Percentage of apolar cells positive or negative for YAP1 during C0 and C1 stages. (C) Percentage of YAP1 positive/negative cells in polar/apolar outer and inner cells of C2 embryos. (D) Percentage of TE and ICM cells of B0-B3 embryos positive/negative for YAP1 and polar/apolar. Brightfield images were taken before fixation. Scale bars, 20µm.

### GATA3 follows YAP1 in outer cells and co-localize in trophectoderm

The establishment of a high-resolution time frame of the appearance of cell polarisation and nuclear YAP1 localisation throughout human pre-implantation development allowed us to match the appearance of YAP1 with GATA3, the first transcription factor associated with TE specification in the human embryo (Gerri et al., 2020).

Confirming our findings shown in Fig 3A-B, the presence of YAP1 positive nuclei was observed at the early compaction stages in 34.1% of cells at C0 and 44.2% at C1 in all embryos (Fig 4A-B). Nuclear GATA3 was first detected at low levels in two out of four compacting C1 embryos in a few cells (11.6%) and always co-localised with nuclear YAP1 (Fig. 4A-C). Throughout compaction, we never observed YAP1-/GATA3+ nuclei. Once distinguishable inner cells were formed at the compacted C2 stage, most of the embryos (n=5/8) showed differential levels of YAP1 and GATA3 between the inner and outer cells. GATA3 was completely absent from inner cells and most inner cells lacked both proteins (75.0%) while 25% were positive for YAP1 only (Fig. 4C; Fig. S1A, C2), as found in previous immunofluorescence analysis (Fig. 3C). At the C2 stage, 26.1% of the outer cells were still negative for both YAP1 and GATA3 (Fig. S1B, C2), 73.9% of the cells showed nuclear YAP1 only and in 21.8% of the cells YAP1 and GATA3 co-localized (Fig. 4C). The differential pattern between inner and outer cells became more pronounced during the transition to the blastocyst stages (B0-B3) (Fig. 4D). Most of the B0 cells were outer cells (83.4%), of which half were YAP1 positive only, 35.9% were YAP1/GATA3 positive (Fig. 4D) and 14.1% lacked either proteins (Fig. S1B). During the following blastocyst stages the percentages of TE cells containing both YAP1 and GATA3 increased: 95.2% in B1, 94.6% in B2 and 97.3 % in B3. Conversely, the percentage of outer/TE cells that contained YAP1 only decreased: 0.8 % in B1, 4.1% in B2 and 0.5% in B3. Furthermore, outer/TE cells that lacked both proteins decreased during blastocyst expansion: 4% in B1, 1.4% in B2 and 2.2% in B3. While the TE cells displayed YAP1/GATA3 in almost all nuclei by the B3 stage, the inner/ICM cells remained negative for both proteins with few exceptions: 87.1% in B0, 93.2% in B1, 89.2% in B2 and 96.2% in B3. Notably, some rare inner/ICM cells were either GATA3 and/or YAP1 positive: B0 (12.9%), B1 (6.8%), B2 (10.7%) and B3 (3.8%) (Fig. 4D; Fig. S1B, B1 panel). The percentage of YAP1 positive cells as well as the nuclear intensity for YAP1 increased between the C0 and C1 stages, and YAP1 appearance preceded GATA3 that became more prominent in C1 (Fig 4B-E). Between C2 and B0, YAP1 and GATA3 became more restricted to the outer cells, with a clear decline in the inner cells. These results confirm reported single cell transcriptomics data (Meistermann *et al*., 2021) (Fig. S2A-B). YAP1 and GATA3 proteins were absent in oocytes (Fig. S3A). We conclude that YAP1 and GATA3 can be found in the nuclei of compacting embryos before polarity is fully established, and that the presence of nuclear YAP1 precedes the appearance of GATA3.

**Figure 4:**
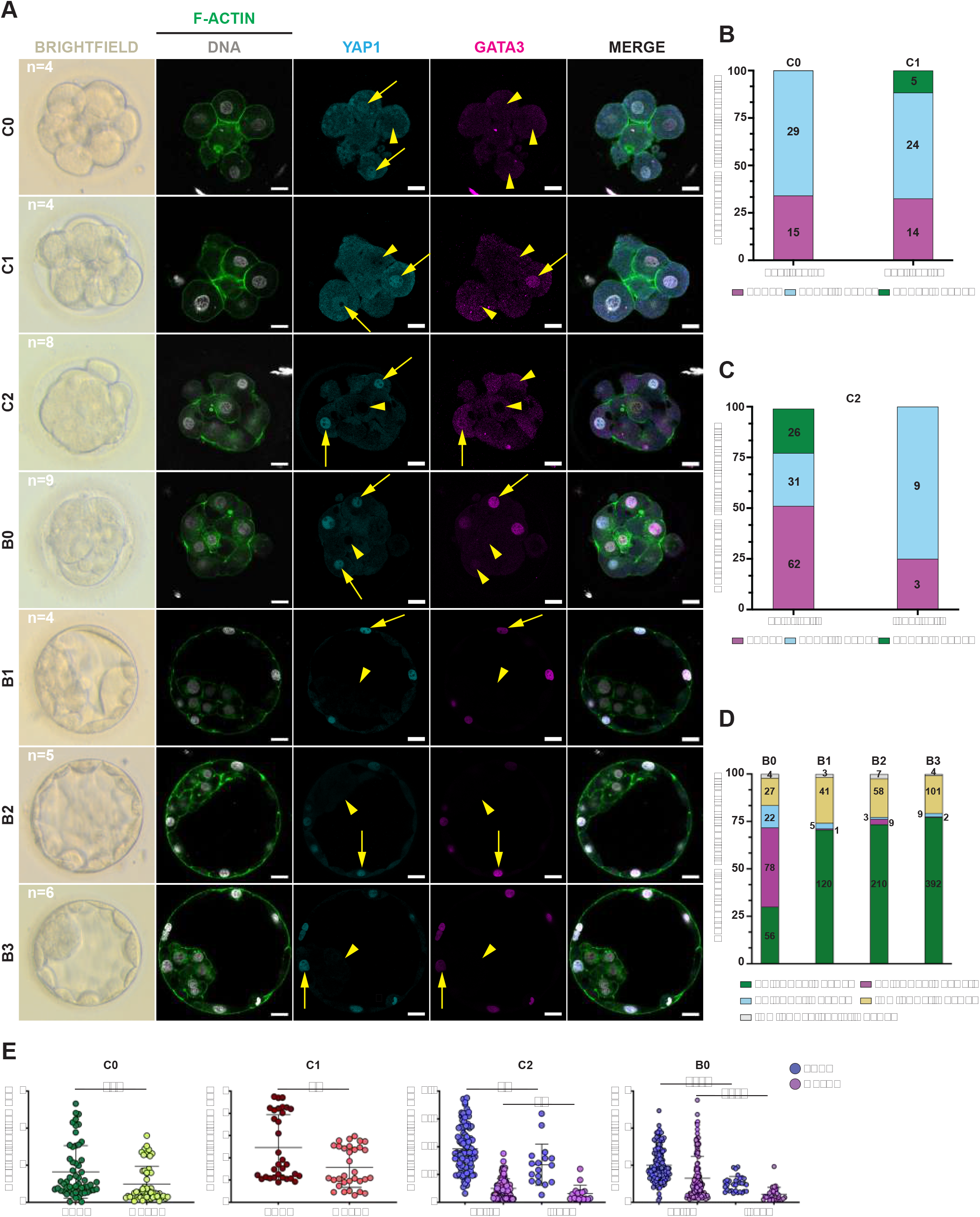
GATA3 and YAP1 co-localize before polarity is established and mainly characterize the outer cells in human pre-implantation embryos. (A) Optical sections of embryos analysed for DNA (white), F-actin (green), YAP1 (turquoise) and GATA3 (magenta), presence indicated by yellow arrows, absence indicated by yellow arrowheads. (B) Percentage of unallocated cells during C0 and C1 stages positive for YAP1, negative for both YAP1 and GATA3 and positive for both YAP1 and GATA3. (C) Percentage of YAP1 and GATA3 positive/negative cells in outer and inner cells of C2 embryos. (D) Percentage of TE and ICM cells of B0-B3 embryos positive or negative for YAP1 and/or GATA3. (E) Nuclear enrichment of YAP1 and GATA3 in C0-B0 embryos. Mann Whitney-*U*-test, **p<0.01, ***p<0.001, ****p<0.0001. Brightfield images were taken before fixation. Scale bars, 20µm.

### TEAD1/YAP1 co-localisation is restricted to (precursor) trophectoderm and primitive endoderm cells

YAP1 is co-activator of the TEAD1-4 family of transcription factors. Recently we showed that in the human embryo TEAD4 proteins are present in all nuclei at the fully compacted embryo stage (Gerri *et al*., 2020). To better understand how TEAD4 is involved in lineage segregation throughout human pre-implantation development, we performed immunofluorescent analysis of TEAD4, p-ERM and GATA3. Low levels of TEAD4 proteins first appeared in all nuclei during compaction (C1) and they were ubiquitously present in the nuclei of all the cells throughout pre-implantation development from the compacted stage onwards (C2-B6) (Fig. 5A-B). This confirmed the single-cell RNAseq data of (Meistermann *et al*., 2021)(Fig S2A-B).

**Figure 5:**
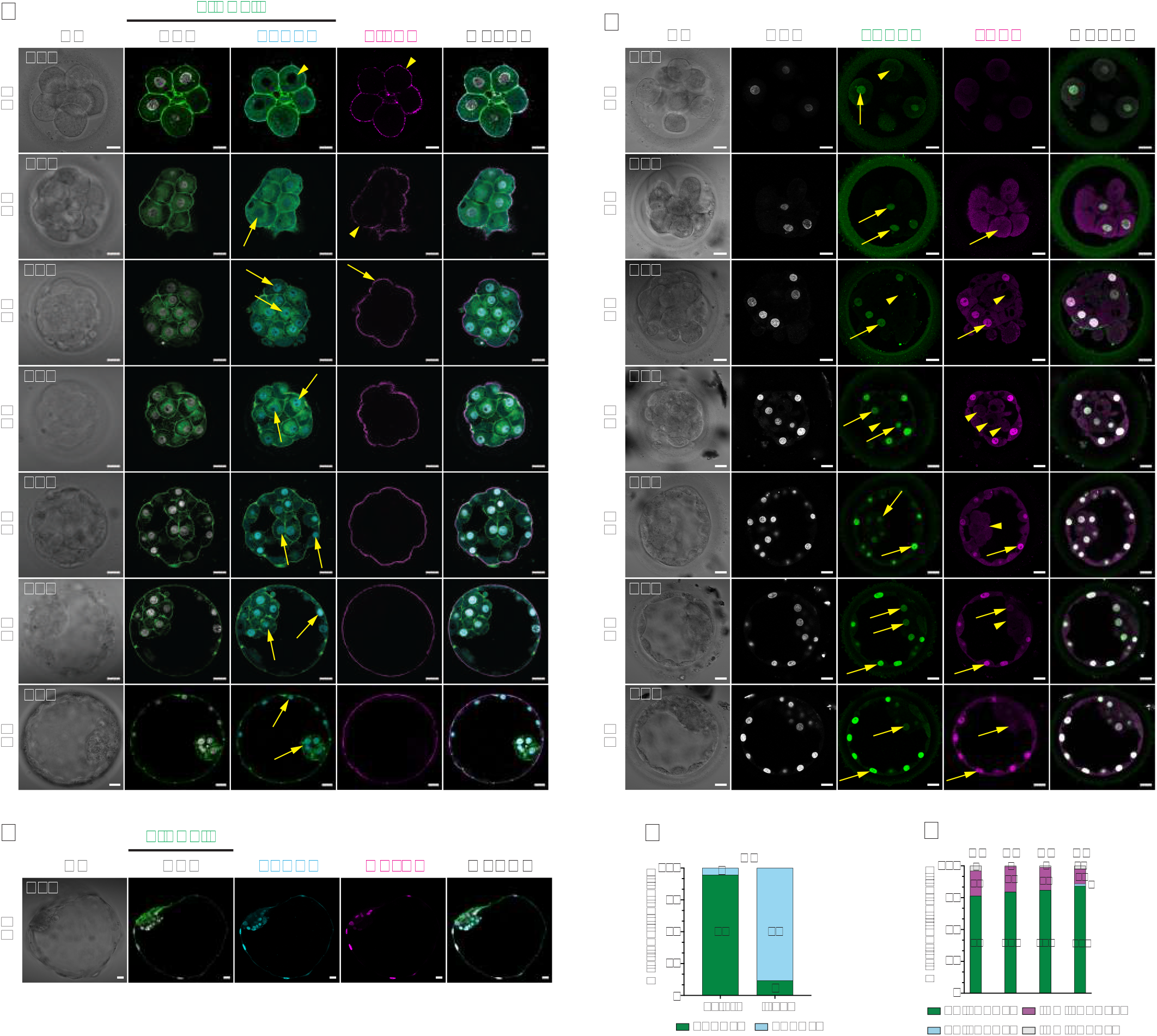
TEAD4 is ubiquitously expressed in the human embryo while TEAD1 consecutively marks the trophectoderm lineage. (A) Optical sections of embryos (C0-B3) analysed for DNA (white), F-actin (green), TEAD4 (turquoise) and p-ERM (magenta). (B) Optical section of a B6 embryo analysed for DNA (white), F-actin (green), TEAD4 (turquoise) and GATA3 (magenta). (C) Optical sections of embryos (C0-B3) analysed for DNA (white), TEAD1 (green), YAP1 (magenta). (D) Percentage of TEAD1 positive/negative cells at the C2 stage. (E) Percentage of TE and ICM cells positive/negative for TEAD1 in B0-B3 stages. Presence of signal indicated by yellow arrows, absence indicated by yellow arrowheads. Brightfield images were taken after fixation. Scale bars, 20µm.

Transcriptomics data suggest that other members of the TEAD family may play a role during human pre-implantation development as well, specifically TEAD1 stood out as a specific regulator of TE and potentially PrE (Yan *et al*., 2013; Stirparo *et al*., 2018; Meistermann *et al*., 2021). We also selected TEAD1 for further study because this protein had already been found to play a role in the second lineage segregation in the mouse (Hashimoto and Sasaki, 2019). TEAD1 displayed a spatial and temporal pattern that was distinct from TEAD4. TEAD1 was found in 65.5% of the nuclei of the cells already at the start of compaction (C0) and in all the cells during compaction (C1) (Fig. 5C, Fig. S4A). From the compacted C2 stage onwards, TEAD1 proteins were abundantly present in the outer and TE cells up to 6 dpf (B0-B4) (Fig. 5C, Fig. S4B, S4C). At the compacted C2 stage, however, only 11.8% of the inner cells were positive for TEAD1. This TEAD1 pattern coincided with YAP1/GATA3 co-localisation in the outer cells (Figure 4A). Most outer/TE cells throughout the B0-B3 blastocyst stages were positive for TEAD1; 100% in B0-B2 and 98.1% in B3. However, TEAD1 proteins were also detected in most nuclei of the inner/ICM cells of the blastocysts on 5 dpf from cavitation onwards, 83.3% in B0, 91.3% in B1, 92.9% in B2 and 83.6% in B3 (Fig. 5C, 5E), but at visibly lower levels as compared to TE cells (Fig. 5C).

By fine-mapping TEAD1, we noticed increased signals of nuclear localisation of this transcription factor in B4 and later blastocysts in cells that could correspond to PrE. In B4 blastocysts, we found TEAD1 positive and negative cells in a heterogeneous pattern (Fig. S4B) and in the cells facing the cavity (Fig. S4C). Strikingly, the increase in the levels of TEAD1 proteins in the ICM cells coincided with the appearance of nuclear YAP1 in these cells (Fig. S4B). In YAP1 positive cells the epiblast marker NANOG was absent (Fig. S4C) but the PrE marker SOX17 was detected (Fig. S4D). Because this pattern is reminiscent of the second lineage segregation, we hypothesised that TEAD1/YAP1 expressing cells in the ICM of expanding blastocysts could be (precursor) PrE-cells.

To test this hypothesis, we first assessed the presence of NANOG during the compaction (C0-C2) and early blastulation stages (B0-B1). After extensive antibody validation (Fig. S4E-G), NANOG was found to be absent at all compaction stages (Fig. S4E). At the B0-B1 blastocyst stages we found NANOG positive and negative nuclei in the ICM cells as well as in the TE cells (Fig. S4F). Furthermore, from the B3 blastocyst stage on, we stained for EPI (NANOG) and PrE (SOX17 and GATA4) markers. In the ICM of B3 blastocysts we found one main population of cells with NANOG+/SOX17-/GATA4-nuclei (89.1%) (Fig. 6A, 6B). The B3 ICM consisted of four additional minor subpopulations: NANOG+/SOX17+/GATA4-cells (5.4%), NANOG-/SOX17-/GATA4-cells (3.9%) and cells in which exceptionally NANOG, SOX17 and GATA4 (0.8%) and GATA4 and SOX17 only co-localized (0.8%) (Fig. 6B, Fig. S5A-D). In seven out of nine B3 blastocysts, nuclear NANOG was found in all the ICM cells (Fig. 6A). Based on all our observations on the second lineage segregation we assessed the average number of cells in the EPI (12.1±6.5 cells) and PrE-lineage (8.4±3.44 cells) at 6dpf (Fig. 6C). At 6 dpf the ICM consisted of two populations only: NANOG+ cells (EPI) and SOX17+/GATA4+ cells (PrE) either in salt-and-pepper (Fig. 6D) or in segregated populations (Fig. 6E). We also detected GATA3 positive cells co-localizing with SOX17 (27%) (Fig. S5E), which were also found in reported single-cell RNAseq data (Meistermann *et al*., 2021) (Fig S2A-B).

**Figure 6:**
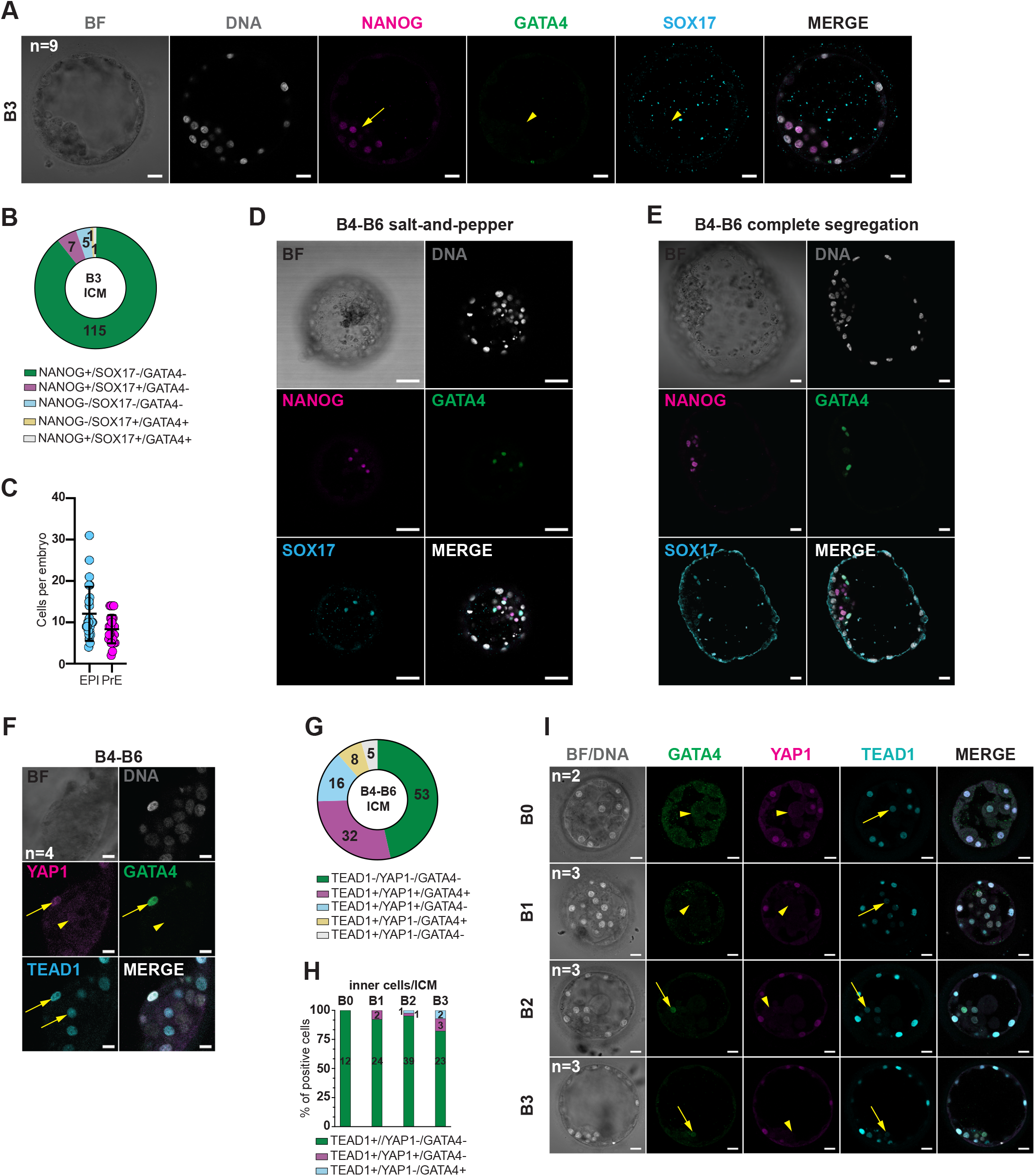
YAP1/TEAD1 consecutively mark primitive endoderm. (A) Optical section of a B3 blastocyst analysed for DNA (white), NANOG (magenta), GATA4 (green), SOX17 (turquoise). (B) Percentage of ICM cells positive/negative for NANOG, GATA4 and SOX17 at the B3 stage. (C) Number of cells per blastocyst in the EPI and PE lineages. Counts are based on NANOG (EPI), SOX17/GATA4 (PE), TEAD1/YAP1 (TE/PE) and GATA3 (TE) protein expression. (D) Optical section of a 6 dpf blastocyst with NANOG, GATA4 and SOX17 in salt-and-pepper constellation. (E) Optical section of a 6 dpf blastocyst with NANOG, GATA4 and SOX17 after completed segregation. (F) Optical section of 6 dpf blastocyst analysed for DNA (white), TEAD1 (turquoise), YAP1 (magenta). (G) Percentage of ICM cells positive/negative for GATA4, TEAD1 and YAP1 at dpf6. (H) Percentage of cells positive/negative for TEAD1, GATA4 and YAP1 in the ICM of B0-B3 blastocysts. (I) Optical sections of B0-B3 blastocysts analysed for DNA (white), GATA4 (green), YAP1 (magenta) and TEAD1 (turquoise). Presence of signal indicated by yellow arrows, absence indicated by yellow arrowheads. Brightfield images were taken after fixation. Scale bars, 20µm.

To investigate the PrE-population further, we co-stained blastocysts for TEAD1, YAP1 and GATA4 (Fig. 6F-I). We identified two main cell populations in the ICM of B4-6 blastocysts (Fig. 6F, 6G): cells negative for all markers that we considered the EPI lineage (46.5%) and cells positive for all three markers that we considered the PrE-lineage (28.1%). We also observed three subpopulations in the ICM cells that consisted of TEAD1+/YAP1+/GATA4-cells (14%), TEAD1+/YAP1-/GATA4+ cells (7%) and TEAD1+/YAP-/GATA4-cells (4.4%). From this pattern it became clear that in the majority of the GATA4 positive PrE-cells co-localisation of TEAD1/YAP1 was found. However, GATA4 was also found in the absence of TEAD1/YAP1 co-localisation (7%). Finally, we analysed embryos from the C0 to the B3 stages for TEAD1, YAP1 and GATA4 (Fig. 6H, 6I, S5F). GATA4 was first detected in ICM cells of B2 (2.4%) and B3 (7.1%) blastocysts when TEAD1 signals were visibly weak and YAP1 was undetectable (Fig. 6H, 6I). Cells positive for all three markers only arise later at B4-6 stage (Fig. 6F, 6G). Therefore, we conclude that TEAD1 and YAP1 co-localize in (precursor) TE and PrE cells.

## Discussion

We show that in human pre-implantation embryos, which compact and polarise heterogeneously between the 8-to 16-cell stages, the formation of a p-ERM apical cap precedes the concentration of F-actin at the apical region. YAP1 and GATA3 co-localize in the polar outer cells, supporting the role of polarity in TE initiation. However, they also co-localize in the nuclei of some blastomeres before the cells are polarized, indicating that the first lineage differentiation event can be initiated independently of polarity. Finally, based on spatial and temporal mapping of transcription factor TEAD1 and co-activator YAP1 we suggest for the first time a role for TEAD1/YAP1 interaction in both TE and PrE differentiation in human pre-implantation embryos.

Compaction and polarisation coincide in human embryos, as in the mouse (Levy *et al*., 1986), but by using morphokinetics it was demonstrated that the human compaction process is heterogeneous (Gerri *et al*., 2020). We confirmed this by use of F-actin staining and we also showed that, as reported in the mouse (Dietrich and Hiiragi, 2007), inner and outer cells are established in the compacted 16-cell stage embryo. We demonstrated polarity by the presence of p-ERM proteins at the apical cortex of the outer cells at the moment of full compaction. Recently Zhu et al. (Zhu *et al*., 2021) revealed that in human embryos polarity occurs in two steps: first F-actin is concentrated at the cell-contact free surface, followed by PAR6/PKC apical cap polarity. Here we demonstrated that this process is preceded by p-ERM polarity, most likely to link F-actin to the plasma membrane, and we suggest that the apical cap in compacted human embryos is established in three steps: first p-ERM, followed by F-actin and finally the PAR complex proteins. Co-localisation of p-ERM, F-actin, PAR6 and aPKC proteins points to maturation of the apical cap when compaction is completed and blastulation starts.

In human embryos, the first lineage differentiation is orchestrated by polarity and YAP1 and GATA3 co-localisation in the compacted embryo (Gerri *et al*., 2020). By carefully examining YAP1/GATA3 co-staining at different stages of development, we observed that YAP1 and GATA3 co-localise in a significant number of nuclei in the compacting/polarising human embryo before inner and outer cells are established and before the apical cap is formed while we never observed GATA3+/YAP1-nuclei throughout compaction. This strongly supports the causal relationship between YAP1 presence and *GATA3* expression (Gerri *et al*., 2020; Zhu *et al*., 2021). We propose, as also suggested by Zhu et al. (Zhu *et al*., 2021), that nuclear YAP1 localisation can be initiated independently of polarity. Nuclear YAP1 could be induced by other mechanisms such as cell-cell contact and/or contractility since YAP1 is very sensitive to mechanical cues (Maître *et al*., 2015; Maître, 2017). We propose that the apical cap reinforces TE initiation, as also suggested by Zhu et al. (Zhu *et al*., 2021). This does not exclude that in polarizing cells nuclear YAP1 is initiated by the polarisation process, which is an ongoing process that is finalized when polarity is fully established, or that nuclear YAP1 is induced by polarity in polarized cells, but this can only be investigated further by live imaging as done in mouse embryos (Gu *et al*., 2022).

In contrast to previous imaging studies in mouse compacted 16-cell stage embryos where non-polarized cells are frequently found on the outside (Anani *et al*., 2014), we did not observe non-polarized cells in an outer position in compacted human embryos. In mouse embryos, these non-polarized cells most likely result from oblique divisions and are actively sorted into an inner position and then assume the ICM fate. We did not detect polarized cells in an inner position in compacted human embryos, as reported in mouse embryos (Anani *et al*., 2014); however, we found YAP1 positive cells in an inner position (25%). Misallocated cells could be sorted to the correct position or could change lineage direction due to plasticity. Cell-sorting events have recently been described in compacting human embryos (Zhu *et al*., 2021). On some occasions, we found YAP1 and GATA3 co-localized in ICM (5 dpf) and PrE (6 dpf) cells, where they most likely originate from miss-positioning during compaction and early blastulation.

In human embryos, YAP1, TEAD1 and TEAD4 are present in the nuclei before compaction and polarity are completely established, as described in the mouse but their role is unknown. YAP1 and TEAD4 proteins were found to play a role in EGA in mouse embryos (Yu *et al*., 2016; Sha *et al*., 2020) and together with TFAP2C and RhoA, TEAD4 was found to play a crucial role in apical domain formation in mice (Zhu *et al*., 2020), but this has not yet been investigated in humans. The upstream signals inducing *YAP1, TEAD1* and *TEAD4* expression in human and mouse embryos are unknown.

We found that TEAD1, which is present in all the blastomeres during compaction, precedes TEAD4. Since TEAD4 is ubiquitously present from the compacted embryos stage onwards while TEAD1/YAP1 are differentially co-localised in what we propose to be (precursor) TE and PrE cells, we suggest that TEAD1 rather than TEAD4 is involved in the segregation of the TE and PrE lineages in humans.

Recently, after *TEAD4* depletion by CRISPR/Cas9 in human zygotes it was found that blastocysts are formed and *CDX2* expression but not *GATA3* expression is downregulated, providing evidence that TEAD4/YAP1 signalling plays a role in the maintenance rather than initiation of TE lineage differentiation (Stamatiadis *et al*., 2022). We suggest that in the compacted human embryo TEAD1/YAP1/GATA3 co-localisation in the outer cells initiates TE lineage differentiation; in inner cells TEAD1, YAP1 and GATA3 do not co-localize and these cells remain pluripotent. This hypothesis needs to be confirmed with functional knock-out studies.

We also demonstrate that the second lineage differentiation event occurs differently in the human and the mouse. In the human embryo, the lineages arising during the second differentiation have been described by immunostainings, but the signalling pathways initiating and maintaining the lineages have not been found yet. GATA6 proteins are present in the nuclei of all the blastomeres after EGA and become restricted in PrE cells when SOX17 and GATA4 are upregulated, in salt-and-pepper pattern and segregation into inner EPI cells and PrE cells facing the cavity (Kimber *et al*., 2008; Kuijk *et al*., 2012; Roode *et al*., 2012; Niakan and Eggan, 2013). While in the mouse embryo, FGF4/FGFR2 signalling plays a key role in the differentiation process, two groups demonstrated that this pathway is not activated in human embryos (Kuijk *et al*., 2012; Roode *et al*., 2012). Here we showed that in human blastocysts the PrE lineage is characterized by the co-localisation of TEAD1 and YAP1. Since the PrE specific markers GATA4 and SOX17 can appear in the absence of TEAD1/YAP1 co-localisation, we hypothesize that the interaction maintains the PrE lineage rather than initiating it. This hypothesis also has to be confirmed with functional studies. This would be different from the mouse model where TEAD/YAP1 interaction orchestrates the expression of *Sox2* (Wicklow *et al*., 2014) and differentiation into the EPI lineage (Hashimoto and Sasaki, 2019). After carefully counting cells expressing NANOG, SOX17 and/or GATA4 in human blastocysts, we found that on 5 dpf in the majority of the B3 blastocysts nuclear NANOG is found in all the ICM cells. In B3 embryos the EPI/PrE-markers co-localize in a small population of the ICM cells (6.2%). On 6 dpf, 2/3 of the ICM cells are EPI cells while 1/3 of the inner cells are PrE cells. According to our data the human TE, EPI and PrE lineages do not emerge simultaneously, as suggested by Petropoulos (Petropoulos, 2016), but in two steps as formulated by Meistermann et al. (Meistermann *et al*., 2021). Our data also support the Meistermann hypothesis (Meistermann *et al*., 2021) that in human blastocysts PrE cells are differentiating from EPI cells in expanding blastocysts (between B3 on 5 dpf and B4 on 6 dpf). The single-cell transcriptomics data (Meistermann *et al*., 2021) and our NANOG analysis also suggest that the ICM (5 dpf) and EPI (6 dpf) are very similar (if not the same), but further characterization of the pluripotent cells is needed to precisely define the stages (Radley *et al*., 2022)

We conclude that TEAD1 and YAP1 co-localize in the (precursor) TE and PrE cells in human pre-implantation embryos. This study establishes a detailed roadmap on polarisation, compaction, position and the presence of lineage specific transcription factors during human pre-implantation development paving the road for further functional studies.

## Author contributions

MR experiments, interpretation, writing, editing, final approval

WE experiments, interpretation, writing, editing, final approval

AD experiments, interpretation, final approval

DD experiments, editing, final approval

LD interpretation, editing, final approval

CG interpretation, editing, final approval

KN interpretation, editing, final approval

GV editing, final approval

HT editing, final approval

JS experiments, editing, final approval

KS interpretation, writing, editing, final approval

HVdV study design, interpretation, writing, editing, final approval

## Acknowledgements

We thank Wilfried Cools from the Biostatistics and Medical Informatics Group (VUB) for the statistical advice.

---

**Supplementary Figure 1:**
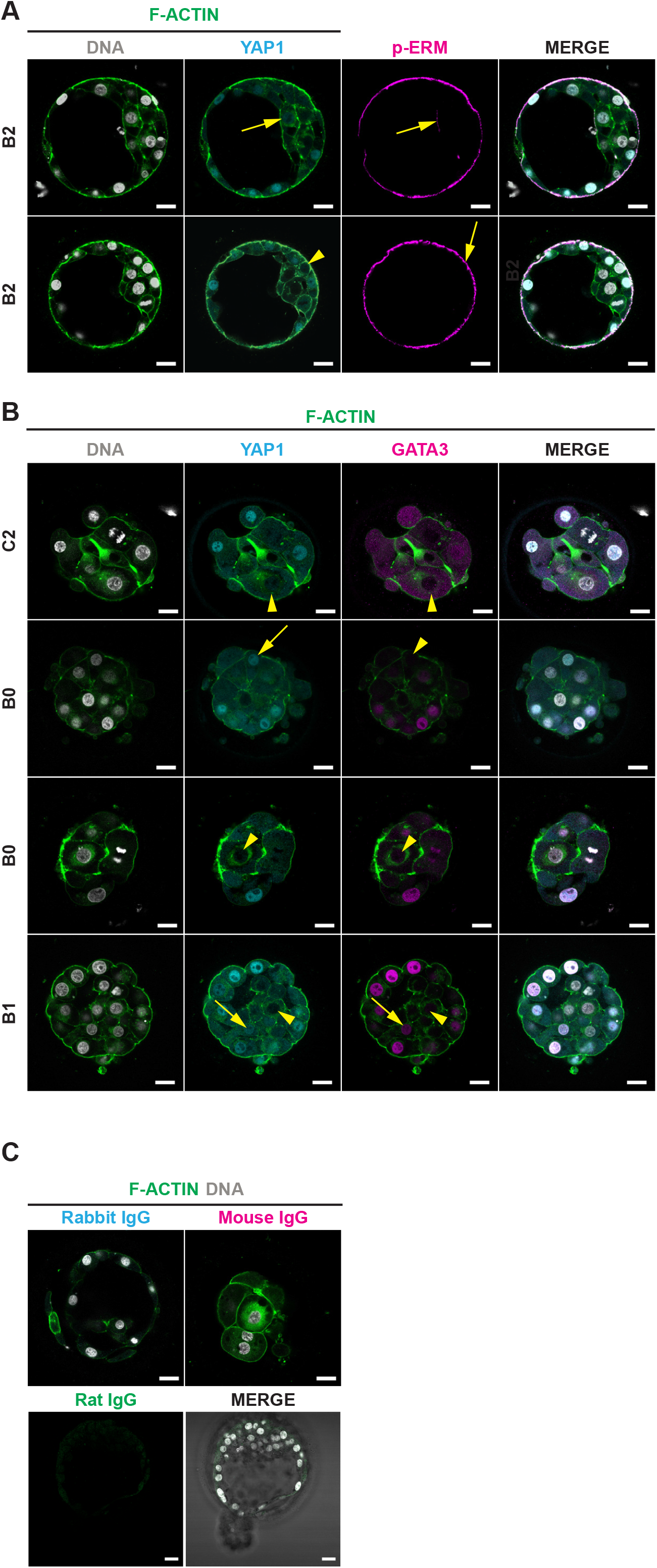
Exceptions of YAP1/p-ERM/GATA3 expression patterns in the human embryo. (A) Optical sections of embryos (B2) analysed for DNA (white), F-actin (green), YAP1 (turquoise) and p-ERM (magenta), presence indicated by yellow arrow, absence indicated by yellow arrowheads. There exist rare cases of polar YAP1-positive cells located in the ICM and YAP1-negative polar TE cells. (B) Optical sections of embryos (C2-B1) analysed for DNA (white), F-actin (green), YAP1 (turquoise) and GATA3 (magenta), presence indicated by yellow arrow, absence indicated by yellow arrowheads. First row shows a C2 stage embryo with a YAP1- and GATA3-negative outer cell. Second row shows a B0 stage embryo with a YAP1-positive, GATA3-negative outer cells. Third row shows a B0 embryos with a YAP1 and GATA3 negative outer cell. Fourth row shows a B1 stage embryo with rare YAP1-positive and GATA3-positive cell in the ICM. (C) Primary antibodies were replaced by either rabbit, mouse or rat IgG as negative control to adjust for possible background in all of our experiments. Brightfield images were taken after fixation. Scale bars, 20µm.

**Supplementary Figure 2:**
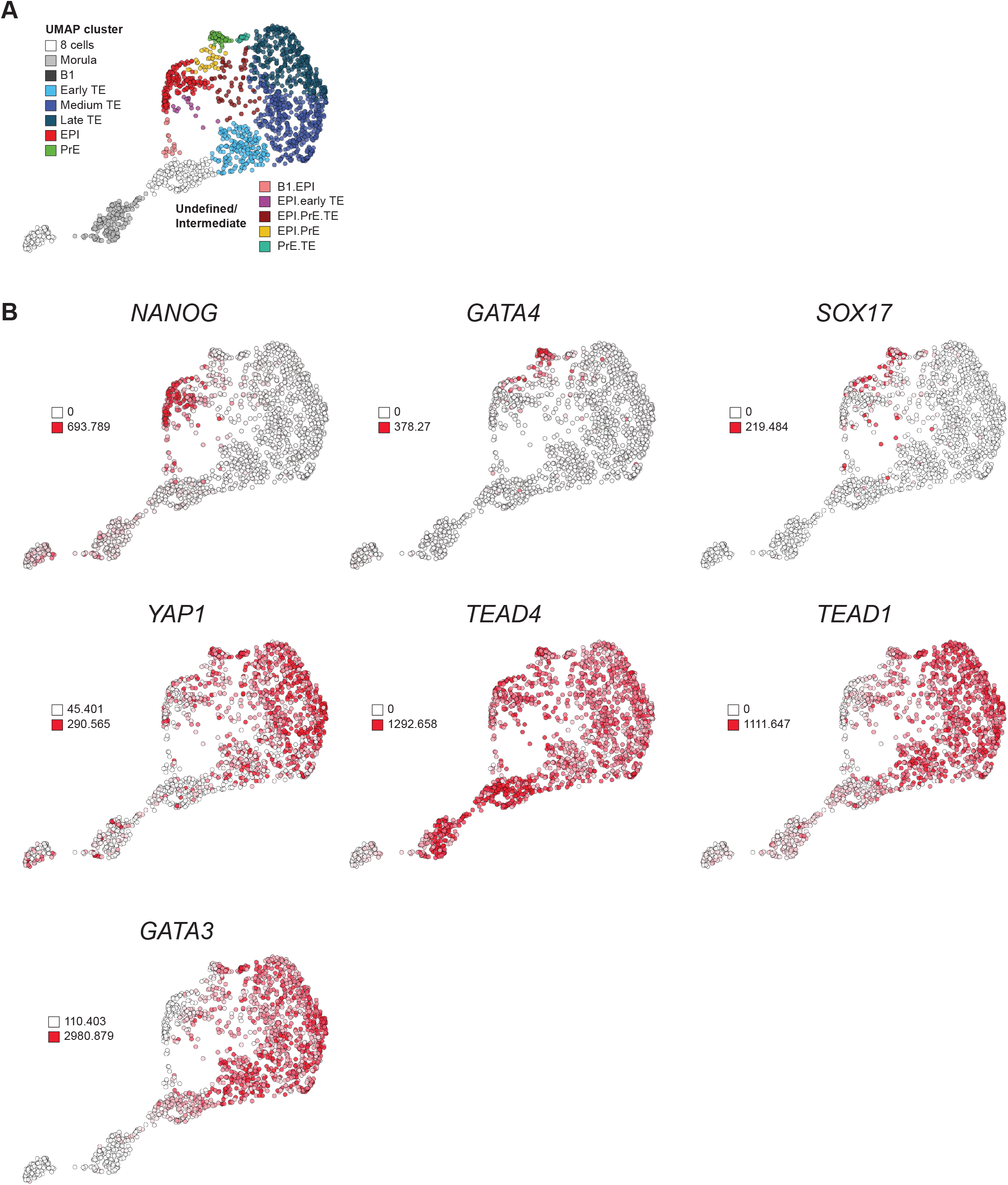
Expression patterns of lineage segregation markers throughout human preimplantation development. (A) Dimension reduction (UMAP) analysis based on (Meistermann *et al*. 2021). The different lineages are indicated by different colours. (B) Projection of lineage marker expression levels of *NANOG, GATA4, SOX17, YAP1, TEAD4, TEAD1* and *GATA3*.

**Supplementary Figure 3:**
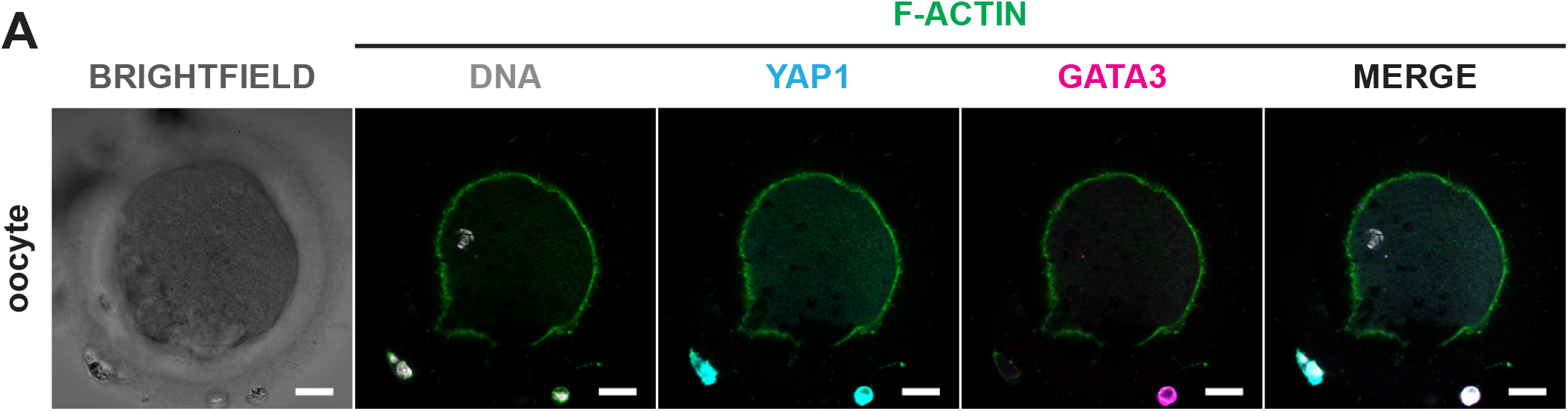
YAP1 and GATA3 are absent in human oocytes. Optical sections of oocytes (n=3) analysed for DNA (white), F-actin (green), YAP1 (turquoise) and GATA3 (magenta), showing absence of both YAP1 and GATA3. Brightfield image was taken after fixation. Scale bars, 20µm.

**Supplementary Figure 4:**
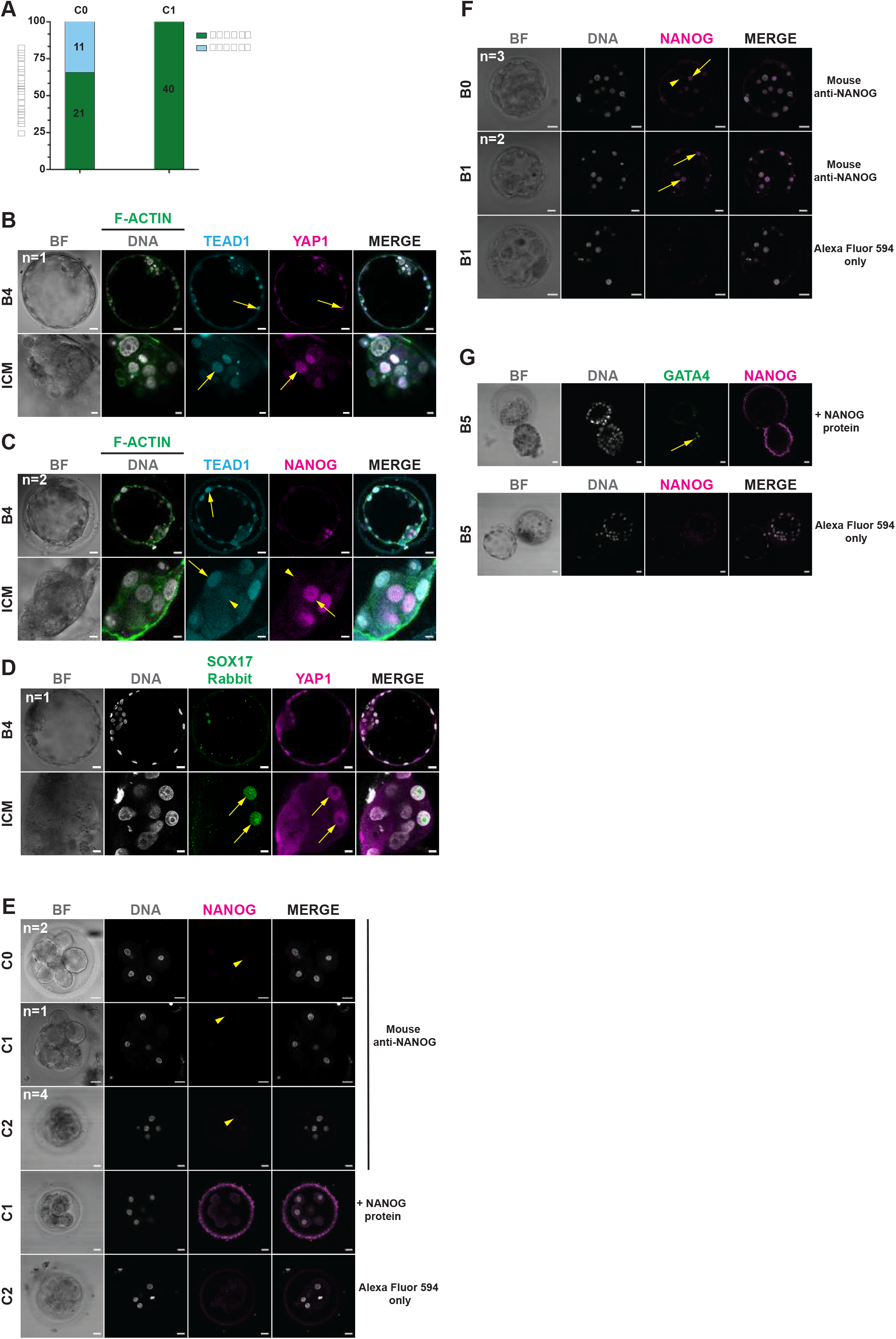
Role of TEAD1 during second lineage segregation and validation of NANOG antibody. (A) Percentage of TEAD1 positive/negative cells at the C0 and C1 stage. (B) Optical section of a B4 embryo analysed for DNA (white), TEAD1 (turquoise) and YAP1 (magenta) with a zoom of the ICM. (C) Optical section of dpf6 embryos analysed for DNA (white), TEAD1 (turquoise), NANOG (magenta). (D) Optical section of a B4 embryo analysed for DNA (white), SOX17 (rabbit, green), and YAP1 (magenta). (E) Validation of the NANOG (magenta) antibody on compaction embryos from C0-C2 stage. First three panels show absence of NANOG using the anti-NANOG antibody. The fourth and fifth panel showed absence of NANOG after adding the NANOG recombinant protein and the secondary antibody only, respectively. (F) Optical sections of B0 and B1 embryos analysed for DNA (white) and NANOG (magenta) after adding the primary antibody (first two panels) or only the secondary antibody (last panel). (G) Optical sections of dpf6 embryos analysed for DNA (white), GATA4 (green) and NANOG (magenta) after adding NANOG recombinant protein (first panel) or the secondary only control for NANOG (second panel). Presence of signal indicated by yellow arrows, absence indicated by yellow arrowheads. Brightfield images were taken after fixation. Scale bars, 20µm.

**Supplementary Figure 5:**
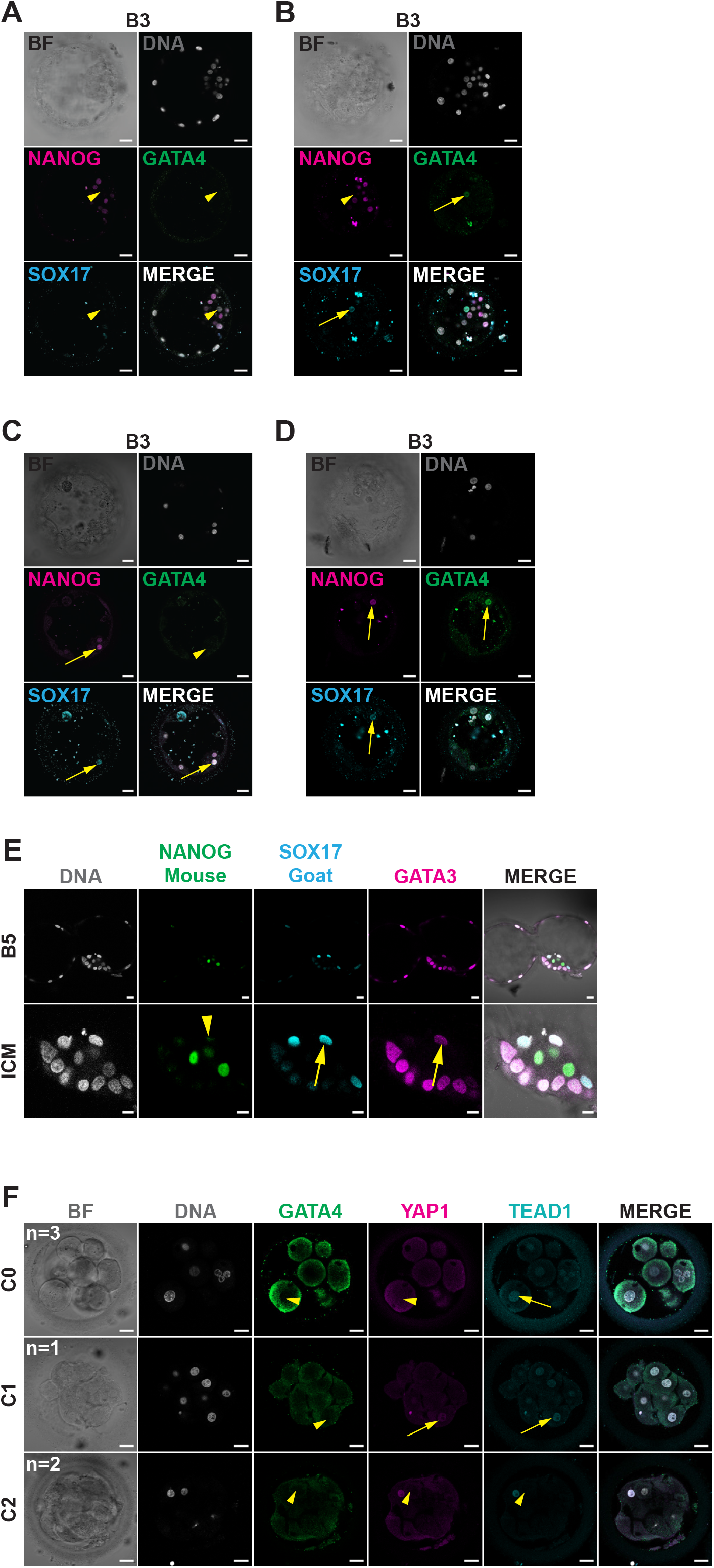
Protein expression of GATA3, NANOG, GATA4 and SOX17 during first and second lineage segregation. (A-D). Optical sections of B3 embryos analysed for DNA (white), NANOG (magenta), GATA4 (green) and SOX17 (turquoise). (E) Optical section of a B5 embryos analysed for NANOG (green), SOX17 (goat, turquoise) and GATA3 (magenta). Note here the presence of a GATA3 positive cell in the PE lineage. (F). Optical sections of C0-C2 embryos analysed for DNA (white), GATA4 (green), YAP1 (magenta) and TEAD1 (turquoise). Presence of signal indicated by yellow arrows, absence indicated by yellow arrowheads. Brightfield images were taken after fixation. Scale bars, 20µm.

