## Supplementary Tables 1 and 2 for "Lineage segregation in human pre-implantation embryos is specified by YAP1 and TEAD1"

**Supplementary Table 1:** Embryo stage scoring used in this study in comparison with the clinical embryologist scoring system from Alpha scientists obtained for human embryos 3 to 6 days post fertilisation (dpf) after imaging confocal microscopy and /or inverted light microscopy.

| Dpf | Confocal/light microscopy | Study score system | Clinical embryologist score system |
| --- | --- | --- | --- |
| 3/4 | Start adhesion | C0 | Cleavage stage |
| 3/4 | Compacting | C1 | C1 |
| 3/4 | Full compaction | C2 | C2 |
| 4/5 | Distinct small cavities | B0 | BL1 |
| 4/5 | 1 small cavity, ICM visible | B1 | BL2 |
| 5 | Full cavity | B2 | BL3 |
| 5 | Expanding | B3 | BL4 |
| Early 6 | Expanded | B4 | BL4 |
| Late 6 | Hatching | B5 | BL5 |
| Late 6 | Fully hatched | B6 | BL6 |

**Supplementary Table 2:** Primary, control and secondary antibodies, F-actin and Hoechst

|  | Reference | Company | Species | Concentration |
| --- | --- | --- | --- | --- |
| YAP1 | H00010413-M01 | Abnova | Mouse Monoclonal | 0.5 µg/mL |
| P-ERM | #3141 | Cell Signalling Technology | Rabbit  Polyclonal | 2 µg/mL |
| GATA4 | **14-9980-82** | Thermofisher | Rat monoclonal | 1µg/mL |
| GATA3 | AB199428 | Abcam | Rabbit  Monoclonal | 2 µg/mL |
| NANOG | AB6273 | Abcam | Mouse monoclonal | 1µg/mL |
| SOX17 | PA5-72815 | Thermofisher | Rabbit polyclonal | 1 µg/mL |
| SOX17 | AF1924 | R&D | Goat polyclonal | 5µg/mL |
| TEAD4 | AB58310 | Abcam | Mouse  monoclonal | 1 µg/mL |
| TEAD1 | 12292S | Cell Signalling Technology | Rabbit monoclonal | 1 µg/mL |
| F-actin | R37110 | Thermofisher | - | 2 drops/mL |
| Mouse Ig | 02-6502 | Thermofisher | - | - |
| Rabbit Ig | sc-2027 | Santa Cruz | - | - |
| Hoechst | H3570 | Thermofisher | - | 5µg/mL |

| **Secondary Antibody** | **Host** | **Company** | **Catalog#** | **Dilution** |
| --- | --- | --- | --- | --- |
| Anti-rabbit IgG, Alexa Fluor 647 | Goat | Invitrogen | A-21245 | 1:200 |
| Anti-mouse IgG, Alexa Fluor 594 | Donkey | Invitrogen | A-21203 | 1:200 |
| Anti-rat IgG, Alexa Fluor 488 | Donkey | Invitrogen | A-21208 | 1:200 |
